# Parallel Likelihood Calculation for Phylogenetic Comparative Models: the SPLITT C++ Library

**DOI:** 10.1101/235739

**Authors:** Venelin Mitov, Tanja Stadler

**Author notes:** Corresponding authors: Venelin Mitov, Department of Biosystem Sciences and Engineering, Swiss Federal Institute of Technology, Mattenstrasse 26, CH-4058 Basel, Switzerland;. Tanja Stadler, Department of Biosystem Sciences and Engineering, Swiss Federal Institute of Technology, Mattenstrasse 26, CH-4058 Basel, Switzerland.

## Abstract

1. Phylogenetic comparative models (PCMs) have been used to study macroevolutionary patterns, to characterize adaptive phenotypic landscapes, to quantify rates of evolution, to measure the heritability of traits, and to test various evolutionary hypotheses. A major obstacle to applying these models has been the complexity of evaluating their likelihood function. Recent works have shown that for many PCMs, the likelihood can be obtained in time proportional to the size of the tree based on post-order tree traversal, also known as *pruning*. Despite this progress, inferring complex multi-trait PCMs on large trees remains a time-intensive task. Here, we study parallelizing the pruning algorithm as a generic technique for speeding-up PCM-inference.
2. We implement several parallel traversal algorithms in the form of a generic C++ library for Serial and Parallel LIneage Traversal of Trees (SPLITT). Based on SPLITT, we provide examples of parallel likelihood evaluation for several popular PCMs, ranging from a single-trait Brownian motion model to complex multi-trait Ornstein-Uhlenbeck and mixed Gaussian phylogenetic models.
3. Using the phylogenetic Ornstein-Uhlenbeck mixed model (POUMM) as a showcase, we run benchmarks on up to 24 CPU cores, reporting up to an order of magnitude parallel speed-up on simulated balanced and unbalanced trees of up to 100,000 tips with up to 16 traits. Noticing that the parallel speed-up depends on multiple factors, the SPLITT library is capable to automatically select the fastest traversal strategy for a given hardware, tree-topology, and data. Combining SPLITT likelihood calculation with adaptive Metropolis sampling on real data, we show that the time for Bayesian POUMM inference on a tree of 10,000 tips can be reduced from several days to minutes.
4. We conclude that parallel pruning effectively accelerates the likelihood calculation and, thus, the statistical inference of Gaussian phylogenetic models. For time-intensive Bayesian inferences, we recommend combining this technique with adaptive Metropolis sampling. Beyond Gaussian models, the parallel tree traversal can be applied to numerous other models, including discrete trait and birth-death population dynamics models. Currently, SPLITT supports multi-core shared memory architectures, but can be extended to distributed memory architectures as well as graphical processing units.

## Introduction

Phylogenetic comparative models (PCMs) have been used for studying the evolution of various biological species, ranging from micro-organisms to animals and plants. Ultimately, these statistical models aim to understand the intricate connections between the macroevolutionary patterns observable in phenotype data from phylogenetically linked species and the fundamental mechanisms of evolution operating on the microevolutionary timescale, such as natural selection and random genetic drift (Lande 1976; Felsenstein 1985; Hansen & Martins 1996; Losos 2011; Uyeda & Harmon 2014; Harmon 2018; Uyeda *et al*. 2018). This quest has led to the recent development of complex multi-trait multi-regime models of evolution (Uyeda & Harmon 2014; Clavel *et al*. 2015; Manceau *et al*. 2016; Bastide *et al*. 2018). The inherent complexity of these models is posing new challenges in terms of parameter inference and model selection.

In their effort to speed-up PCM inference, recent works have shown that, for a broad family of PCMs, the likelihood of an observed phylogenetic tree and data conditioned on the model parameters can be computed in time proportional to the size of the tree (FitzJohn 2012; Ho & Ané 2014; Goolsby *et al*. 2016; Mitov *et al*. 2018). This family includes Gaussian models like Brownian motion (BM) and Ornstein-Uhlenbeck (OU) phylogenetic models as well as some non-Gaussian models like phylogenetic logistic regression (Paradis & Claude 2002; Ives & Garland 2010; Ho & Ané 2014; Mitov *et al*. 2018). All of these likelihood calculation techniques rely on post-order tree traversal also known as *pruning* (Felsenstein 1973, 1981, 1983).

For moderate numbers of traits, combining pruning algorithms for likelihood calculation with gradient-based optimization (Boyd & Vandenberghe 2004) enables maximum likelihood model inference within seconds on contemporary computers, even for phylogenies of many thousands of tips (Ho & Ané 2014). Despite its simple interpretation and several useful statistical properties, the maximum likelihood estimator (MLE) has often been criticised for being a point estimator, uninformative about the likelihood surface, often prone to be a local optimum, and failing to quantify the uncertainty of *a priori* assumed models for comparative data (Bishop 2007; Uyeda & Harmon 2014).

As an elegant alternative, Bayesian approaches such as Markov Chain Monte Carlo (MCMC) allow incorporating prior biological knowledge in the model inference, provide posterior samples and high posterior density (HPD) intervals for the model parameters and, in the case of multi-regime models, integrate the inference of shifts in the evolutionary regimes driven by the dynamics of the adaptive phenotypic landscape (Slater *et al*. 2012a; FitzJohn 2012; Uyeda & Harmon 2014). In contrast with ML inference, though, Bayesian inference methods require many orders of magnitude more likelihood evaluations. This presents a bottleneck in Bayesian analysis, in particular, for complex models of many unknown parameters or when faced with large phylogenies of many thousands of tips, such as transmission trees from large-scale epidemiological studies, e.g. Alizon *et al*. (2010); Shirreff *et al*. (2013); Hodcroft *et al*. (2014); Bertels *et al*. (2017); Mitov & Stadler (2018). While big data should provide the needed statistical power to fit a complex model, the time needed to perform a full scale Bayesian fit often limits the choice to a faster but less informative ML-inference, or a Bayesian inference of a simplified model.

Speeding-up Bayesian inference is an active topic in applied statistics with recent advances that can be classified in several groups. One group of methods are adaptive variants of the random walk Metropolis (RWM) algorithm (Metropolis *et al*. 1953) that aim to decrease the number of MCMC iterations by performing “on-the-fly” changes of the jump distribution, based on what has been “learned” about the parameter space from past iterations (Haario *et al*. 2001; Vihola 2012). A major advantage of these methods is that they are generic with respect to the models and can be implemented as general purpose Metropolis samplers (e.g. `adaptMCMC` (Scheidegger 2017)). A second group are “pre-fetching” methods which modify the Metropolis-Hastings algorithm so that it speculatively executes sequences of individual likelihood calls in parallel, “hoping” that these sequences tend to match the actual accepted states of the MCMC (Brockwell 2006; Angelino *et al*. 2014). Another possibility to use multiple processor power, which could potentially be combined with the above methods, is to delegate the parallelization problem to a low level linear algebra library, e.g. OpenBLAS (Wang *et al*. 2013).

This article contributes to a separate body of work, namely, the ensemble of model-specific approaches that parallelize the likelihood calculation by using specific features of the likelihood function. These include factorizations of the likelihood into a product of components associated with conditionally independent subsets of the model parameters (Whiley & Wilson 2004; Goudie *et al*. 2017) or the observed variables (Ayres *et al*. 2012). Often, this factorization relies on strong model assumptions, such as a hierarchical structure of the model parameters or independence of the observed variables. A common approach used in software packages like BEAST (Drummond *et al*. 2012; Bouckaert *et al*. 2014) is to combine the factorization with caching and reusing of some of the previously calculated likelihood components in consecutive MCMC iterations, as long as these are not affected by the proposed jump in parameter space.

For a phylogenetic comparative model, though, the likelihood cannot (in general) be factorized across parameter groups, trait independence is acceptable only as a null hypothesis and, with a moderate number of traits and pruning likelihood calculation, parallelizing algebraic operations (on low-dimensional vectors and matrices) is inefficient. Hence, we explore the parallelization of the likelihood calculation at the level of traversing the phylogenetic tree, that is, the pruning itself. Parallel tree traversal has been studied in computer science, mostly for the purposes of parallel tree contraction (Reif 1989), automated task scheduling (Qamnieh 2015) and for phylogenetic inference from multiple sequence alignment data (Ayres *et al*. 2012; Ayres & Cummings 2017). Capitalizing on the same ideas, we developed **SPLITT**: a shared-memory C++ library for Serial and Parallel Lineage Traversal of Trees. While we focus on Gaussian phylogenetic models as the main application of the library, we designed the SPLITT programming interface to be generic with respect to the node-traversal operations, hoping that the library could potentially find use in different models, including birth-death population models and discrete trait models. We tested SPLITT on large trees (up to N=100,000) and on different topologies, including balanced and highly unbalanced trees. These tests proved a nice property of the parallel pruning algorithm, namely the fact that its parallel efficiency increases with the tree size as well as the complexity of the node-traversal operations. Thus, for large trees and complex models, the parallel speed-up is limited either by the number of available processors or by another limited resource such as the memory bandwidth. Finally, we showcase that our parallel pruning algorithm coupled with adaptive Metropolis samplers dramatically reduces the time for Bayesian analysis of trees with thousands of tips.

## Materials and Methods

In this section, we introduce SPLITT and show through an example how it is used to parallelize a pruning algorithm over a given phylogenetic tree and data. Further technical details and examples are provided in Sections 2 and 3 of the Supplementary Information and the SPLITT online documentation.

### A general framework for parallel tree traversal

SPLITT implements a general framework for specifying the type of trait data, the model parameters and the “node-traversal” operations, which are executed in a pre-order or a post-order traversal of the tree (Section 1, Supplementary Information). The node-traversal operations represent user-defined rules specifying how a set of variables associated with each node, called a “node-state”, is initialized and updated in the computer memory, based on the input tree and data, the model parameters, and the node-states of the previously visited nodes. At the end of the traversal, the final node-state values are accessible for calculating a quantity of interest, such as the likelihood of model, given the tree and the data.

The node-states can be calculated in parallel for any group of siblings or more remote cousins on the tree. Formally, SPLITT makes the following key assumption:

##### Assumption 1

Calculating the state of a node *j* can be done independently from the calculation of the state of any other node *k*, provided that neither *j* is an ancestor of *k*, nor *k* is an ancestor of *j*.

To maximize the potential for parallel execution, the lifecycle of a node during traversal is divided in three operations (Fig. 1c,d):

1. *InitNode*: initializes the node-state based on the input data and model parameters only. This operation does not depend on the states of other nodes. Hence, it is fully parallelizable.
2. *VisitNode*: “updates” the state of its operand node based on the state of either the node’s parent if in pre-order traversal, or the node’s daughters if in post-order traversal. This operation can be executed in parallel for any group of nodes satisfying Assumption 1 and having their parents *Visit*-ed (if a pre-order traversal) or their daughters *Prune*-ed (if a post-order traversal, see PruneNode below). To prevent a possible race-condition, in a post-order traversal, this operation should not modify the state of the parent or any ancestor of the operand node. Executing this operation on the root node is optional and not done by default.
3. *PruneNode* (post-order traversal only): “communicates” the state of a node to its parent node. SPLITT ensures that this operation is synchronized between siblings, i.e. daughters of the same parent node. Hence, this operation is convenient for accumulation (e.g. summation) of state-variables of the daughters into the state of their parent (Fig. 1c). This operation is not defined for the root of the tree.

The parallel speed-up can depend on multiple factors, including the balancedness of the tree, and the computing and memory complexity of the traversal operations, which can be different between nodes in the tree. Noticing that there is no *one-size-fit-all* parallel traversal strategy that guarantees fastest execution, previous works have studied queue-based and range-based parallelization strategies (Reif 1989; Qamnieh 2015; Ayres & Cummings 2017). SPLITT implements such algorithms called “orders” (Section 1.1, Supplementary Information). As a default setting, SPLITT implements a mode “auto”, in which it compares the execution time of different parallel orders during the first several calls on a given tree and data, choosing the fastest one for all subsequent calls.

**Figure 1:**
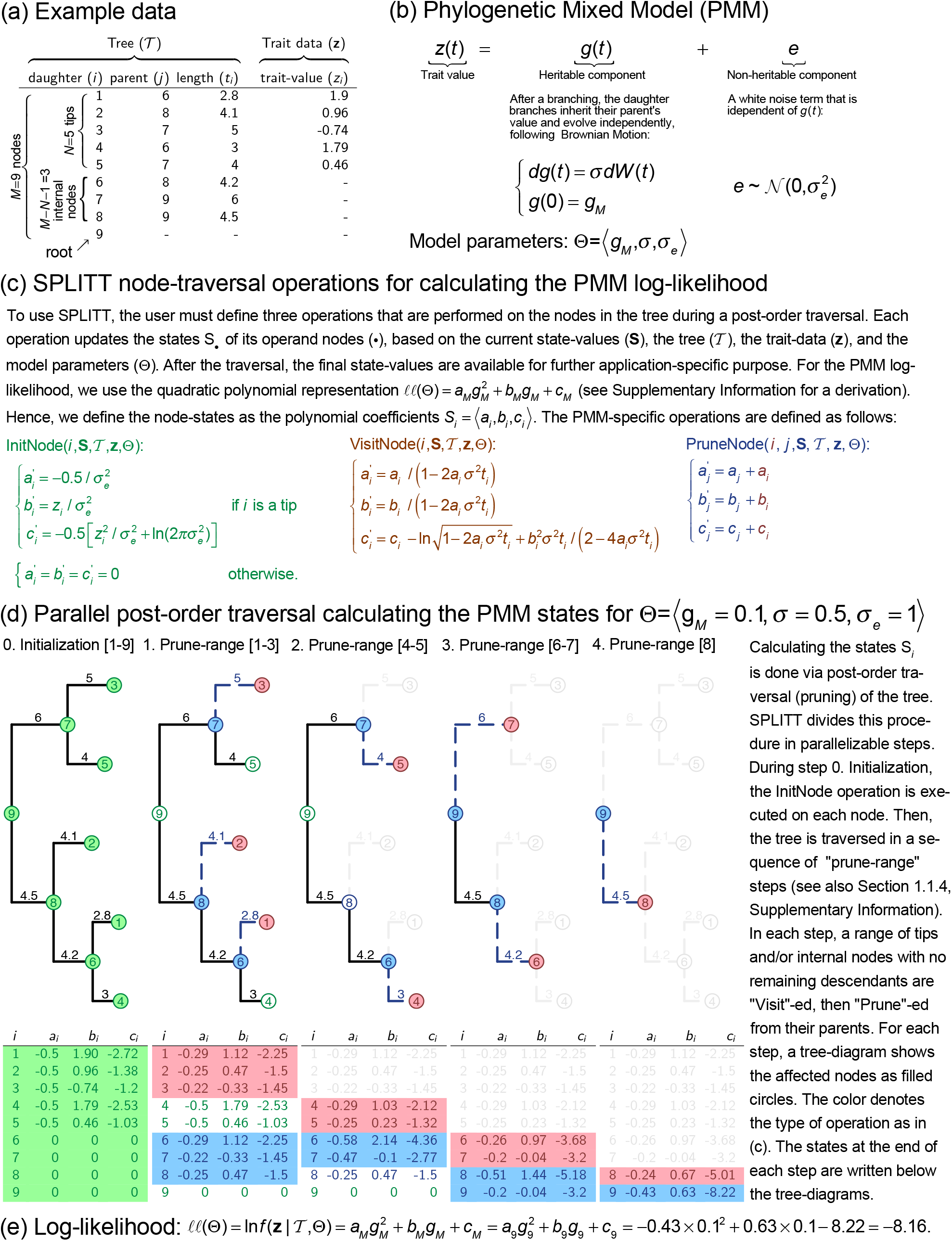
Parallel pruning for calculating the log-likelihood of the phylogenetic mixed model.

### A showcase: the phylogenetic (Ornstein-Uhlenbeck) mixed model

To illustrate the use and to test the SPLITT library, we developed two variants of the so called phylogenetic mixed model (PMM) – the original PMM assuming a Brownian motion process (Lynch 1991; Housworth *et al*. 2004), and its recent extension to an Ornstein-Uhlenbeck process, the POUMM, which we and other authors have used previously to analyze the evolution of set-point viral load in HIV patients (Mitov & Stadler (2018) and references therein).

Figure 1b shows the mathematical formulation of the PMM (see Section 2.2, Supplementary Information for a more general mathematical formulation of the P(OU)MM model and a biological interpretation of its parameters).

The key assumption enabling a pruning algorithm to evaluate the P(OU)MM likelihood is that the trait evolves independently in each lineage descending from a branching point in *𝓣*. This allows to factorize the likelihood function over the sub-trees in *𝓣*, treating the values of *g* at the branching points as unknown variables which are integrated over their distributions expected under the model parameters, Θ. This integration leads to a simple formulation of the P(OU)MM log-likelihood as a quadratic polynomial of *g_M_* (Theorem S1, Section 2.2, Supplementary Information):

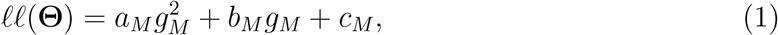

where the coefficients *a_M_, b_M_* and *c_M_* are functions of the tree topology, branch lengths, the observed trait values and the model parameters. Denoting by *Desc(j)* the set of direct descendants (daughters) of node *j*, for the PMM, these functions are given by the following recursive formulas (Theorem S1, Section 2.2, Supplementary Information):

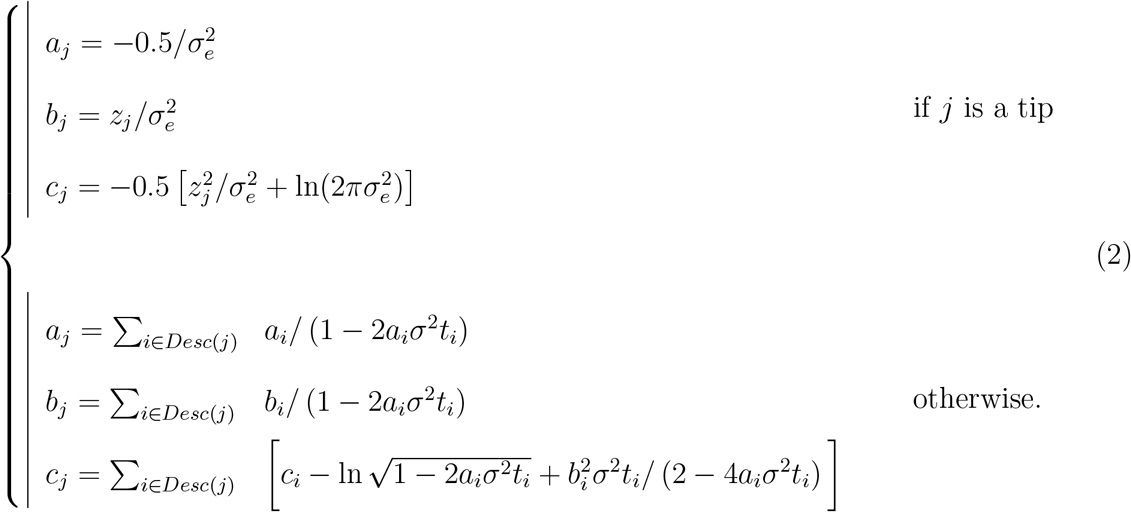

Based on Eq. 2, we define the node-traversal operations (InitNode, VisitNode and PruneNode) as shown on Fig. 1c. Fig. 1d shows how the node-states are initialized and updated in parallel “prune-ranges” for an example tree and trait data and model parameters (Fig. 1a,d). After the traversal the values *a_M_, b_M_* and *c_M_* from the root-state are plugged in Eq. 1 to obtain the log-likelihood value (Fig. 1e).

### Generalization to multi-trait Ornstein-Uhlenbeck and mixed Gaussian phylogenetic models

The quadratic polynomial representation of the log-likelihood function (Eq. 1) can be generalized to a broader family of models. In Section 2.2, Supplementary Information, we show how the coefficients *a_M_, b_M_* and *c_M_* can be calculated for the single-trait Ornstein-Uhlenbeck mixed model. In a separate work (Mitov *et al*. 2018), we extend this integration technique to evaluate the likelihood of multi-trait Ornstein-Uhlenbeck and mixed Gaussian phylogenetic models, i.e. models in which different types of models assigned to different lineages of the tree. These models have been implemented in several R-packages summarized in the following sub-section.

### Other pruning algorithm examples

In Sections 2.1 and 2.2.2, Supplementary Information, we describe another pruning algorithm for calculating the POUMM log-likelihood, which is based on the generalized 3-point structure algorithm (Ho & Ané 2014). In Section 2.3, Supplementary Information, we give an example of a pruning algorithm for calculating the likelihood of a discrete (binary) observed at the tips of a phylogenetic tree.

### Software

We provide SPLITT as a C++ library licensed under version 3.0 of the GNU Lesser General Public License (LGPL v3.0) and available on https://github.com/venelin/SPLITT.git. In its current implementation, the library uses the C++11 language standard, the standard template library (STL) and the OpenMP standard for parallel processing.

The single-trait POUMM has been implemented in the R-package POUMM, available on https://github.com/venelin/POUMM.git. Section 3, Supplementary Information provides details on the model inference procedure implemented within the package and reports a test of technical correctness.

The generalization to a multi-trait mixed Gaussian phylogenetic model (MGPM) has been implemented in the R-package PCMBase (Mitov *et al*. 2018) available on https://github.com/venelin/PCMBase.git. An accompanying package called PCMBaseCpp, which is based on SPLITT, a provides a parallel C++ implementation of the likelihood calculation for the MGPM model. This package is available at https://github.com/venelin/PCMBaseCpp.git.

### Technical correctness

To test the technical correctness of the SPLITT library and the higher level POUMM, PCMBase and PCMBaseCpp packages, we used the method of posterior quantiles (Cook *et al*. 2006). For the single-trait POUMM implementation (POUMM R-package), we report the technical correctness test in Section 3.2, Supplementary Information. For the multi-trait implementation (PCMBase and PCMBaseCpp R-packages), the technical validation is reported in Mitov *et al*. (2018).

## Results

We evaluated the performance of the SPLITT library using the single-trait and multi-trait Phylogenetic Ornstein-Uhlenbeck Mixed Model (POUMM) as a showcase. The single-trait POUMM was implemented in the R-package POUMM (Section 3, Supplementary Information), based on the quadratic polynomial representation of the log-likelihood (Section 2.2, Supplementary Information). The multi-trait POUMM version was implemented in the R-package, `PCMBaseCpp`, using a multi-trait generalization of the quadratic polynomial representation described in Mitov *et al*. (2018). The POUMM is a suitable model for a comparative benchmark, because a number of R-packages provide similar OU-based phylogenetic models, using C++ for the likelihood implementation. These include, among others, `geiger` (Pennell *et al*. 2014) and `diversitree` (FitzJohn 2012) for the single-trait case and `Rphylopars` (Goolsby *et al*. 2016) for the multi-trait case.

We used the R-package `apTreeshape` (Bortolussi *et al*. 2012) to generate tree topologies of sizes *N ∈* {100; 1000; 10, 000; 100, 000}. To generate the random trees, we used the function `rtreeshape()` with a biased model. A parameter *p* in this model controls the disproportion of branching rates for the left and right lineages starting from a given parent node. For each *N*, we used four settings for *p* as follows:

1. *p* = 0.5 corresponding to equal left and right branching rates and resulting in balanced trees;
2. *p* = 0.1 corresponding to unbalanced trees in which one of any two sibling branches (sharing the same parent node) splits at rate *p* = 0.1, while the other splits at rate *p’* = 1 – *p* = 0.9 (time units are arbitrary, so we can assume that the rates correspond to splitting probabilities per unit time).
3. *p* = 0.01 corresponding to very unbalanced trees (splitting rates of *p* = 0.01 and *p’* = 0.99 for any couple of sibling branches);
4. *p* = 0.01/N corresponding to a ladder-like tree (see Fig. 2).

This resulted in a total of 16 topologies (trees for *N* = 1, 000 shown on Fig. 2). For each topology, random branch lengths were assigned overwriting the default branch lengths of 1 assigned by `rtreeshape()`. Since the OU-implementations in the current `diversitree` and `Rphylopars` versions do not support non-ultrametric trees, each tree was ultrametrized (adjusting branch lengths so that all tips have the same root-tip distance). For each tree, we generated random trait-values by simulating under the POUMM model using random parameters.

**Figure 2:**
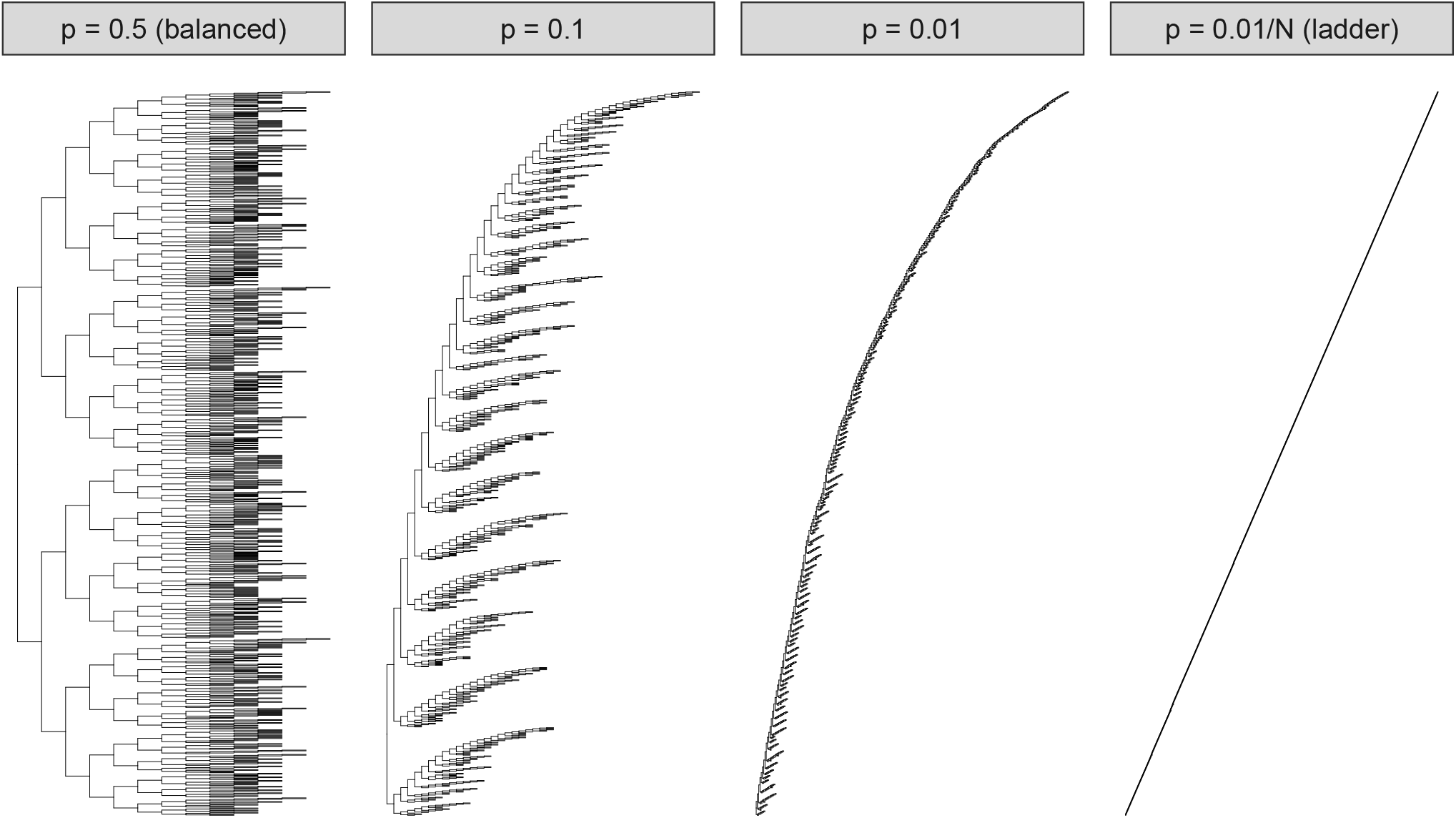
Test tree topologies for *N* = 1, 000. For visualization purpose, all branch lengths have been set to 1, whereas the random branch lengths were used in the benchmarks. Note that the tree for *p* = 0.5 is nearly but not perfectly balanced due to the random nature of the tree generation process, as well as *N* not being an exact degree of 2.

### Time for preprocessing the tree

Each of the tested packages implements a preprocessing step initializing cached data-structures that are re-used during likelihood calculation. In the case of `SPLITT`, this is the constructor-function of the internal `Tree` structure; in the case of `diversitree`, this is the function `make.ou`; in the case of `geiger`, this is the internal function `bm.lik`. We note that the time for creating the cache structure is not important in scenarios of fitting Gaussian phylogenetic models to a fixed tree and data (created once, at the beginning of the inference process). However, these times become important in the case when the tree topology is inferred together with the model parameters from trait and sequence alignment data.

We measured the preprocessing time on the 16 trees (Table 1). The times scaled linearly with the size of the tree for the packages using the SPLITT library (`POUMM` and `PCMBaseCpp`) and for `diversitree`. For these packages the time was not affected by the unbalancedness of the tree. For `geiger`, we observed longer times, both for bigger *N* as well as for more unbalanced trees. For *N* = 100, 000 and *p* = 0.01/`N`, both, `diversitree` and `geiger` failed with a `stack-overflow` error. The relatively short times for the SPLITT-based `POUMM` and `PCMBaseCpp` packages indicate that `SPLITT` could potentially be used for phylogenetic inference.

### Time for POUMM likelihood calculation

To measure the likelihood calculation time, we ran performance benchmarks on a personal computer (PC) running OS X on an Intel(R) Core(TM) i7-4850HQ CPU @ 2.30GHz with 4 CPU cores, and on the “Euler” scientific cluster (https://scicomp.ethz.ch/wiki/Euler) running Linux OS on an Intel(R) Xeon(R) CPU E5-2680 v3 @ 2.50GHz running 24 physical cores. Here, we comment on the calculation times on the PC, noting that the times on Euler for up to 4 CPU cores were nearly equal (Figs. S2-S6, Supplementary Information).

**Table 1.** Times for tree-preprocessing in milliseconds.

| N | Implementation | p=0.5 | p=0.1 | p=0.01 | p=0.01/N |
| --- | --- | --- | --- | --- | --- |
| 100 | geiger | 5 | 6 | 9 | 9 |
| 100 | diversitree | 4 | 4 | 4 | 4 |
| 100 | SPLITT | 2 | 2 | 2 | 1 |
| 1,000 | geiger | 18 | 26 | 78 | 414 |
| 1,000 | diversitree | 20 | 20 | 22 | 30 |
| 1,000 | SPLITT | 3 | 2 | 3 | 3 |
| 10,000 | geiger | 358 | 449 | 1,345 | 355,396 |
| 10,000 | diversitree | 207 | 211 | 227 | 1,338 |
| 10,000 | SPLITT | 14 | 13 | 13 | 15 |
| 100,000 | geiger | 20,215 | 21,629 | 36,349 | - |
| 100,000 | diversitree | 2,421 | 2,619 | 2,883 | - |
| 100,000 | SPLITT | 130 | 131 | 131 | 140 |

We distinguish the different implementations according to the following criteria:

- Number of traits: we distinguish between single-trait implementations, i.e. `geiger`, `diversitree` and `POUMM`, and multi-trait implementations, i.e. `Rphylopars` and `PCMBaseCpp`. For the multi-trait implementations, we measured the time for 1, 4, 8 and 16 traits.
- Mode: denotes whether the implementation is single threaded using one physical core of the CPU – serial, or multi-threaded, running as many threads as there are physical CPU cores – parallel;
- Order: denotes the order in which the prune-able nodes are processed. We tested three possible orders: postorder – the nodes are processed sequentially; queue-based – the nodes are processed in parallel as they enter the queue (Algorithm S1, Section 2, Supplementary Information), synchronized thread access to the queue; range-based – the nodes in each pruning generation are processed in order of their allocation in memory, no need for a synchronized access to a queue (Algorithm S2, Section 2, Supplementary Information).
- Implementation: the R-package and the back-end used (R or C++).

The likelihood calculation time was measured using the R-function “sys.time” calling the specific likelihood implementation on a fixed set of parameters *n* = 100 times, then, dividing the cumulative time by n. To avoid influence from other processes running on the same PC, the benchmark was run after a restart of the operating system (OS). The resulting times for the single-trait implementations running on the PC are shown on Fig. 3.

On small trees of 100 tips, the fastest single-trait POUMM implementations were the serial C++ implementations from the packages POUMM and diversitree (about 0.03 ms); the range-based parallel implementation was nearly as fast on balanced trees (*p* = 0.5) but was progressively slower on unbalanced trees. The geiger implementation was nearly an order of magnitude slower (0.2 ms). The POUMM queue-based parallel implementation was nearly 100 times slower (nearly 2 ms), presumably due to the excessive synchronization overhead. The serial R implementation from the diversitree package was the slowest (above 2 ms), which was expected, since the R interpreter is notorious for its slow speed compared to compiled languages like C++. On bigger balanced trees (*N* > 100, *p* = 0.5), the range-based parallel implementation took over, reaching up to 4× speed-up with respect to the range-based serial implementation, up to 5× speed-up with respect to the postorder serial implementation and up to 10× speed-up with respect to the `diversitree` serial C++ implementation. This reveals a consistent speed-up for all trees except the ladder tree, where parallelization of the internal nodes is not possible (see Fig. 1b and 2). The time for the other serial implementations and the POUMM queue-based parallel implementation scaled up linearly with *N*.

The times for the multi-trait implementations running on the PC are shown on Fig. 4. For these implementations, the likelihood calculation times were about two orders of magnitude higher compared to the single-trait implementations. This is due to slow algebraic operations, for example arithmetic division in the single-trait case as opposed to matrix inversion in the multi-trait case.

**Figure 3:**
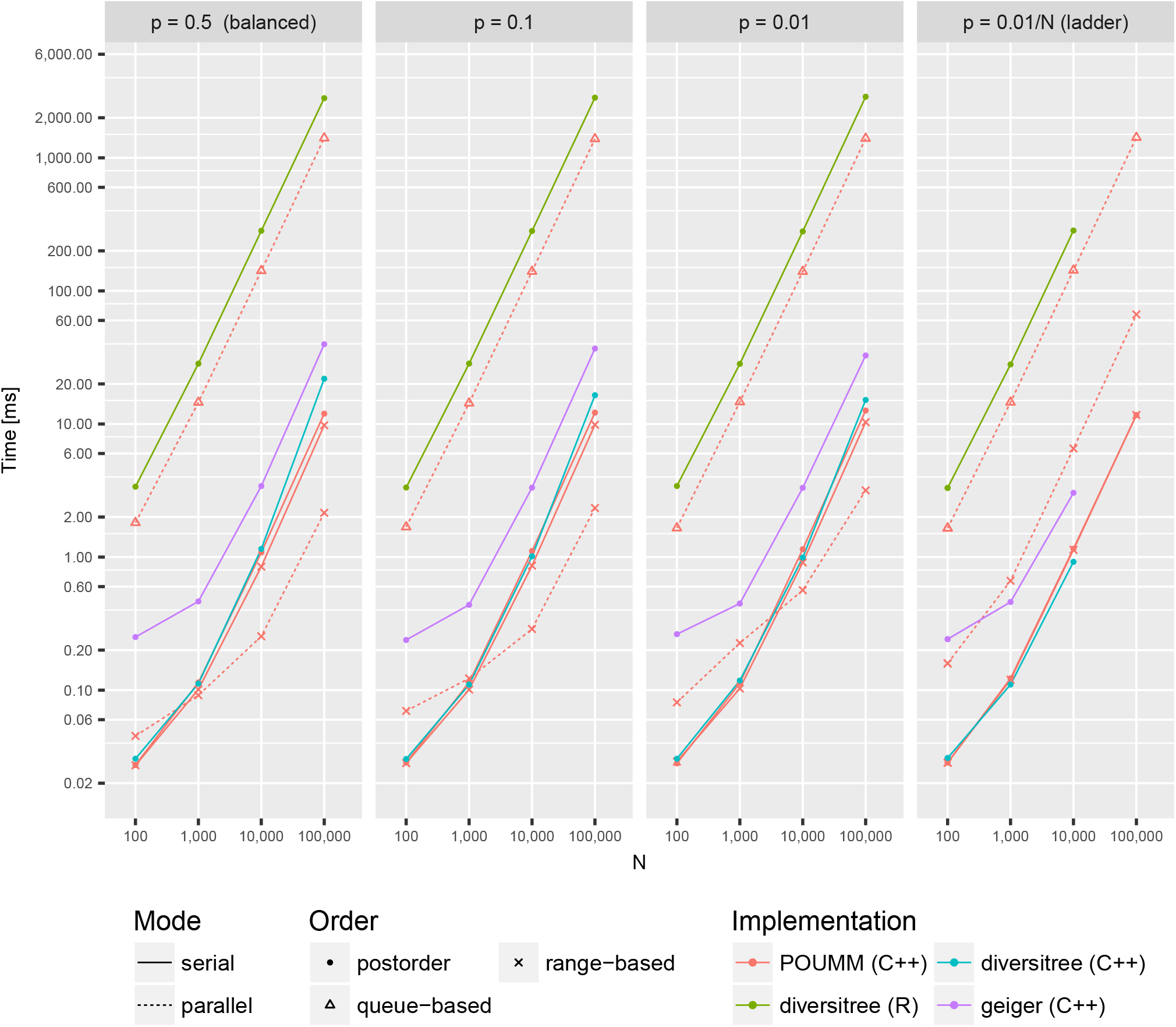
Likelihood calculation times for single-trait R and C++ implementations of the POUMM model on a PC (processor Intel(R) Core(TM) i7-4850HQ CPU @ 2.30GHz with four physical cores). Both, the *x*-axis denoting the number of tips in the tree and the *y*-axis denoting the calculation time in milliseconds are on a log-10 scale. Panels from left to right correspond to different tree topologies with left-most panel corresponding to a perfectly balanced tree and right-most panel corresponding to a ladder tree, see also Fig. 2.

### Parallel speedup

The parallel speed-ups for the Euler cluster benchmark for single-trait implementations and for multi-trait implementations with 16 traits are shown on figs. 5 and 6 (see also Figs. S7-S9, for multi-trait implementations with 1, 4 and 8 traits).

For single-trait implementations, the parallel speed-up is negligible for trees of less than 1000 tips and for highly unbalanced trees (Fig. 5). The parallel speed-up becomes noticeable for large balanced trees, peaking at 10x for a balanced tree of 100,000 tips, running on 20 CPU cores (Fig. 5). The above behaviour is explained by the fact that the InitNode and VisitNode operations in the single-trait case are very fast relative to the thread-management operations. Also noteworthy is the fact that even on balanced trees above 100,000 tips, the parallel efficiency, i.e. the ratio of the parallel speed-up and the number of parallel cores, drops below 50% when running on more than 20 CPU cores. This suggests a possible competition between the CPU cores for a limited resource such as the processor cache or the memory bandwidth.

For the multi-trait implementations, the InitNode and VisitNode operations are computationally more intensive. This is why we observe substantial parallel speed-up on the smallest as well as the most unbalanced trees trees (Fig. 6). However, for all multi-trait cases, we observe a decline in parallel speed-up with more than 12 CPU cores (Fig. 6). The most reasonable explanation for this is competition between the CPU cores for a limited hardware resource.

### Combined parallel likelihood calculation with adaptive Metropolis sampling

In Bayesian MCMC inference, the parallel likelihood calculation can be combined with an adaptive MCMC sampler. The POUMM R-package implements this approach by embedding the SPLIT-based likelihood calculation in a Metropolis sampler with coerced acceptance rate available from the `adaptMCMC` R-package (Scheidegger 2017) (Section 3.1, Supplementary Information). We tested this approach during a POUMM analysis of a transmission tree from the HIV epidemic in UK (N=8483) reported elsewhere (Mitov & Stadler 2018). This showed faster MCMC convergence (Fig. S10, Section 5, Supplementary Materials) and overall less than 30 minutes for the MCMC run (Section 5, Supplementary Materials).

**Figure 4:**
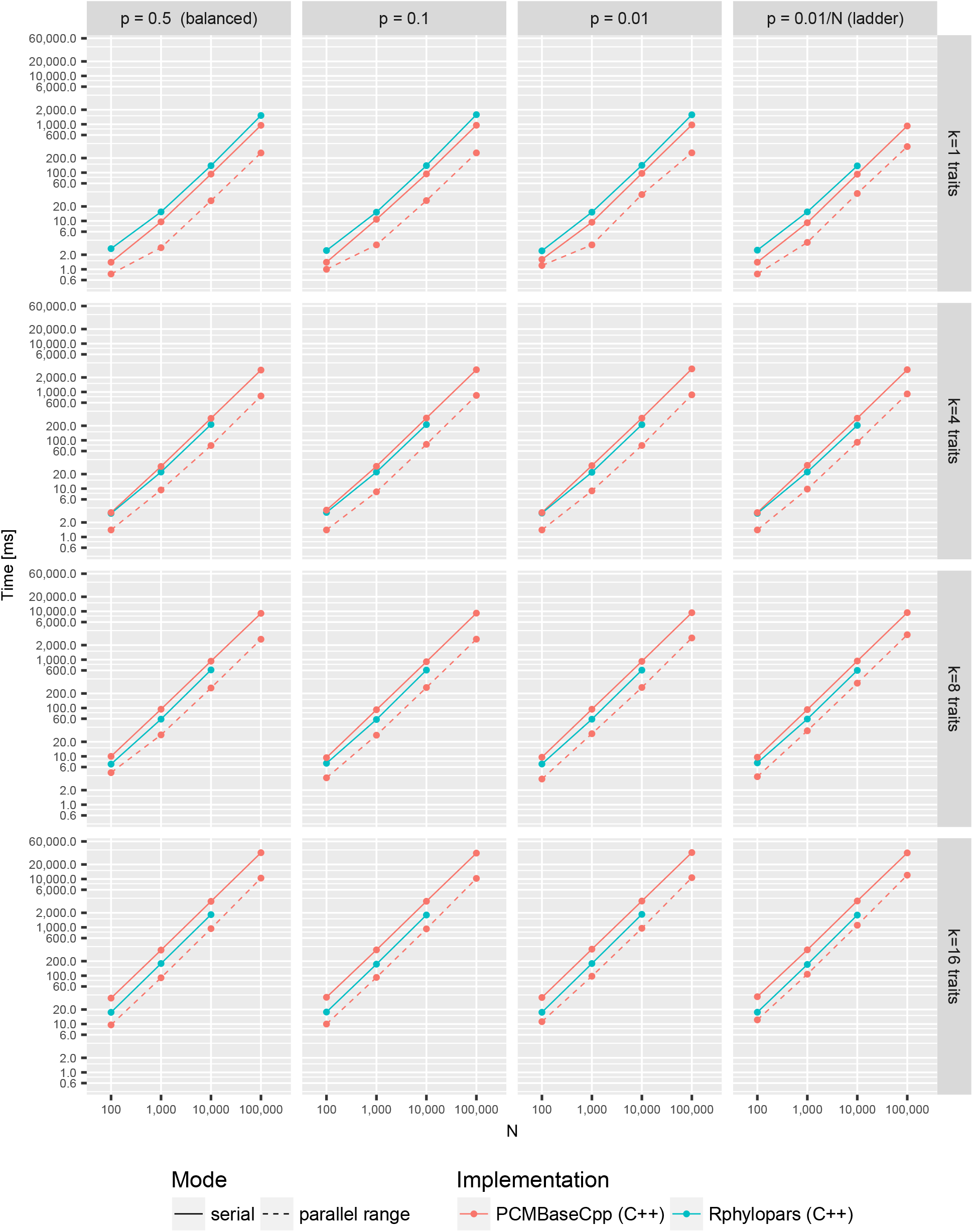
Likelihood calculation times for multi-trait C++ implementations of the POUMM model on a personal computer (processor Intel(R) Core(TM) i7-4850HQ CPU @ 2.30GHz with four physical cores). For simplicity, only serial and parallel range modes are shown, noting that the parallel queue mode had slightly slower times compared to the parallel range mode.

**Figure 5:**
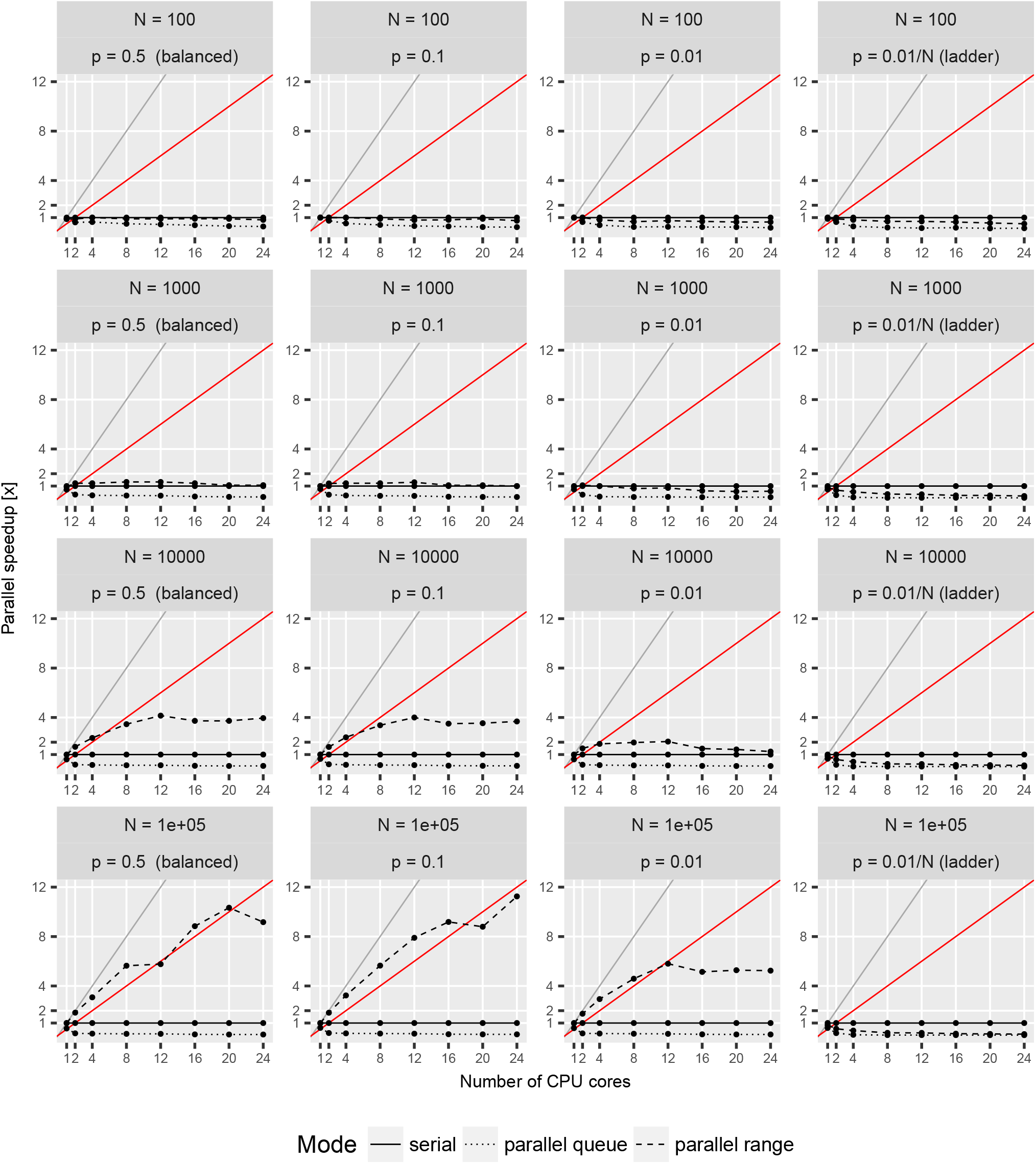
Parallel speed-up for the single-trait POUMM implementation on the Euler cluster (package `POUMM`). The grey and red lines denote the expected speed-up at 100% and 50% parallel efficiency, respectively. Horizontally, the panels correspond to the different tree topologies, see also Fig. 2. Vertically, the panels correspond to the different tree-sizes.

**Figure 6:**
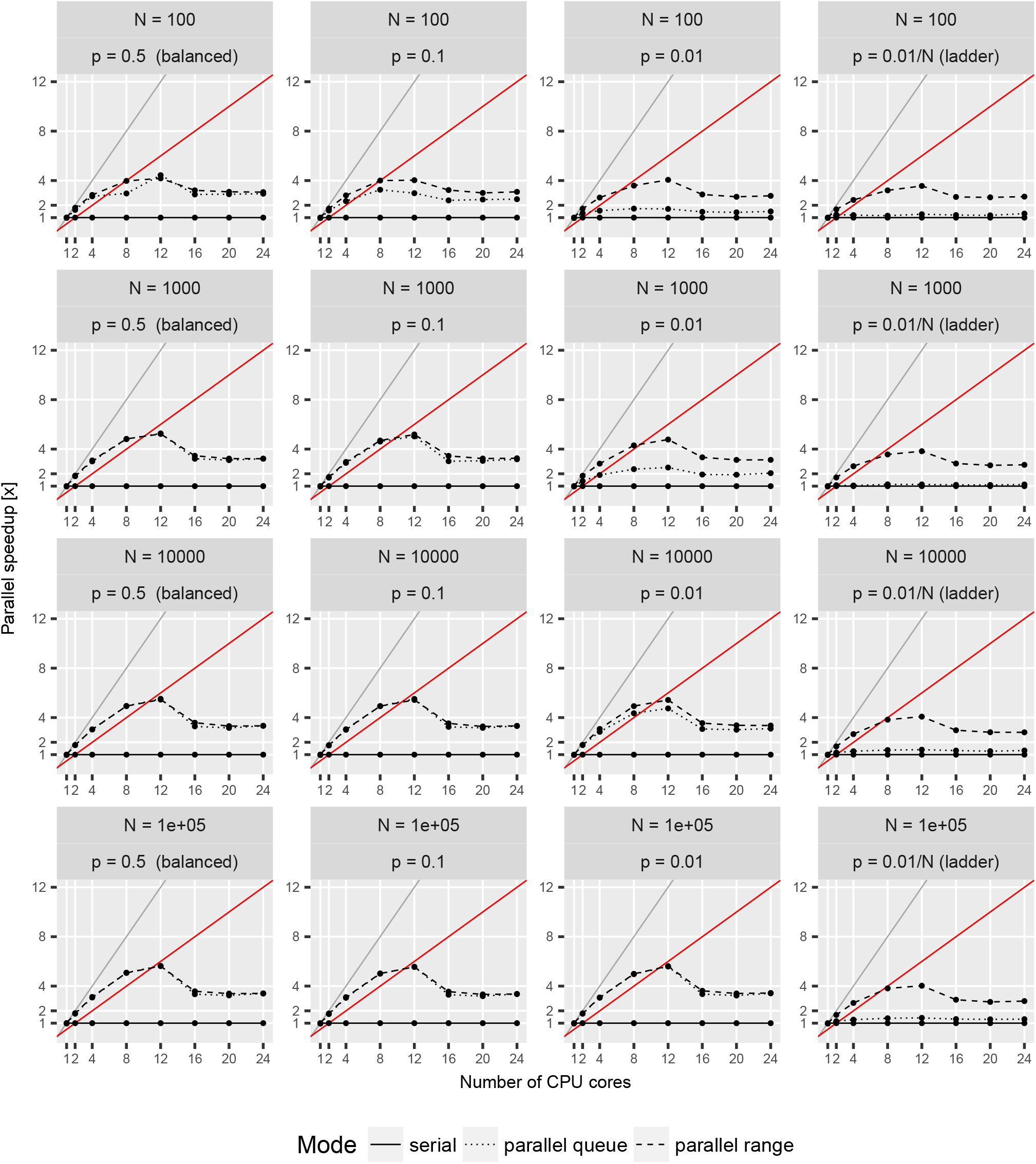
Parallel speed-up for the multi-trait (*k* = 16 traits) POUMM implementation (package `PCMBaseCpp`) on the Euler cluster. The grey and red lines denote the expected speed-up at 100% and 50% parallel efficiency, respectively. Horizontally, the panels correspond to the different tree topologies, see also Fig. 2. Vertically, the panels correspond to the different tree-sizes.

## Discussion

The examples in this article focused on Gaussian models of continuous trait evolution (see Materials and Methods and Section 2, Supplementary Information). Yet, SPLITT can in principle be used for any algorithm that runs a pre-order or post-order tree traversal. For example, another family of models where SPLITT could be used are models of structured populations. When calculating the likelihood for a phylogenetic tree under a structured birth-death model, the calculations proceed in a pruning fashion (Kühnert *et al*. 2016) and may be improved with respect to speed using our approach. However, the structured coalescent likelihood for a tree is a function of all co-existing lineages even for approximate methods (Müller *et al*. 2017), and thus a pruning formulation is not available.

We did not develop examples of pre-order traversal. One such example is the simulation of traits evolving along the tree, which can be used for validation and approximate inference of phylogenetic models. In complex phylogenetic comparative models, where an exact calculation of the likelihood is elusive or computationally intractable, it is possible to use simulations of trait evolution along the tree for approximate likelihood calculation (Kutsukake & Innan 2013) or approximate Bayesian computation (ABC) (Slater *et al*. 2012b). Both approaches are computationally intensive and could benefit from parallel execution using SPLITT.

We should not omit mentioning other software libraries implementing parallel likelihood computation of different Markov models of sequence evolution. For example, several high level tools for ML and Bayesian tree inference, e.g. Drummond *et al*. (2012); Bouckaert *et al*. (2014); Ronquist & Huelsenbeck (2003), use the library BEAGLE which distributes the computation for the independent sites of the sequence alignment among multiple CPU or GPU cores (Ayres *et al*. 2012). SPLITT operates on a different level, namely, it parallelizes the computation for independent lineages in the tree. Both approaches are interesting because they fit well to different sizes of the input data – while BEAGLE achieves significant parallel speed-ups in long alignments comprising many thousands nucleotide or codon columns (Ayres *et al*. 2012), SPLITT is better suited to shorter alignments of potentially many thousands of species.

Based on the performance benchmarks, we conclude that with the current implementation of SPLITT, running on the above-mentioned hardware, the parallel speed-up from parallel tree traversal is up to one order of magnitude using up to 20 CPU cores. A future GPU-based extension of SPLITT would show if it can reach higher levels of parallel speed-up and efficiency. Reaching higher speed-up of the Bayesian inference, though, is possible if the parallel traversal likelihood calculation is combined with a general purpose adaptive Metropolis sample. An example application of this combined approach to real data is given in Section 5 in Supplementary Information.

### Outlook

The past decade has seen a rapid advance in the production of multi-core processors. At the same time, it appears that the maximum clock frequency of a single processing unit is approaching the maximum achievable for semi-conductor based architectures. In parallel with this development on the hardware side, the volume of sequence data and the size of phylogenetic trees is growing exponentially. For instance, in less than five years the size of phylogenetic trees used for calculating the heritability of HIV virulence has increased from a few hundreds to several thousand patients (Alizon *et al*. 2010; Hodcroft *et al*. 2014). This motivates the development of novel parallel algorithms capitalizing on the multi-core technology. The parallel tree traversal library, SPLITT, enables parallel computation for a vast set of phylogenetic models, facing the challenges of increasing model complexity and volumes of data in phylogenetic analysis.

## Supplementary Material

Technical details, supplementary figures and supplementary tables are provided in Supplementary Information. Data from the performance benchmarks and simulations for technical correctness is accessible from the SPLITT github page https://github.com/venelin/SPLITT. The `POUMM` package and user guide is available at https://github.com/venelin/POUMM. The `PCMBaseCpp` package is available at https://github.com/venelin/PCMBaseCpp.

## Acknowledgements

We thank Dr. Krzysztof Bartoszek for valuable insights on the Ornstein-Uhlenbeck process.

## Author Contributions Statement

VM conceived the ideas and designed the methodology; VM implemented the software; VM and TS planned the performance benchmarks and the technical correctness tests; VM led the writing of the manuscript. Both authors contributed critically to the drafts and gave final approval for publication.

